# TraB is a novel component of the ER-Mitochondria contact site (EMCS) with dual roles in ER-Mitochondrial tethering and mitophagy

**DOI:** 10.1101/2022.02.25.481886

**Authors:** Chengyang Li, Patrick Duckney, Tong Zhang, Yanshu Fu, Xin Li, Johan Kroon, Geert De Jaeger, Yunjiang Cheng, Patrick J Hussey, Pengwei Wang

**Affiliations:** Key Laboratory of Horticultural Plant Biology (MOE), Huazhong Agricultural University, Wuhan, 430070, China; Hubei Hongshan Laboratory, Wuhan, 430070, China; Department of Biosciences, Durham University, South Road, Durham, DH1 3LE, UK; Department of Plant Biotechnology and Bioinformatics, Ghent University, Ghent, Belgium

**Keywords:** Membrane contact sties, Endoplasmic Reticulum, Mitochondria, Autophagy, Mitophagy, Arabidopsis

## Abstract

ER-mitochondria contact sites (EMCSs) are important for mitochondrial function. Here, we have identified a novel eukaryotic EMCS complex, comprising a family of uncharacterised mitochondrial outer-membrane proteins, TraB1 and the ER protein, VAP27-1. In Arabidopsis, there are two TraB1 isoforms and the *trab1a/trab1b* double mutant exhibits abnormal mitochondrial morphology, strong starch accumulation and impaired energy metabolism, indicating that these proteins are essential for normal mitochondrial function. Moreover, TraB1 proteins also interact with ATG8 in order to regulate mitochondrial degradation (mitophagy). The turnover of depolarised mitochondria is significantly reduced in both *trab1a/b* and VAP27 mutants (*vap27-1/3/4/6)* under mitochondrial stress conditions, with an increased population of dysfunctional mitochondria present in the cytoplasm. Consequently, plant recovery after stress is significantly perturbed. A similar phenotype is found in both autophagy mutants (*atg5* and *atg7*), suggesting that TraB1 regulated mitophagy and ER-mitochondrial tethering are two closely related processes, necessary for normal mitochondrial function. Taken together, we ascribe a dual role to TraB1 which is a novel component of the EMCS complex in eukaryotes, regulating both tethering of the mitochondria to the ER and mitophagy.

## Introduction

In plant cells, organelles interact with each other to regulate multiple physiological activities^1,2,3,4,5^. Such intimate contact between the different organelles generates an interactive network which facilitates rapid material and signal exchange. The endoplasmic reticulum (ER) is physically connected to other membrane structures through membrane contact sites^6^ (MCS), including the ER-PM contact sites^7,8^ (EPCSs) and ER-mitochondria contact sites^4,9^ (ERMCs). The ERMC interactions in particular are essential in multiple physiological activities, for example Ca^2+^ transfer, lipid metabolism, autophagy, mitochondrial dynamics and morphogenesis^3,4,9,10,11,12^. However, the protein components and biological relevance of EMCSs are poorly understood in plants compared to their counterparts in animals and fungi.

In order to maintain cellular homeostasis, impaired mitochondria have to be removed effectively through a process called mitophagy, which is a highly selective autophagic process. The successful removal of damaged mitochondria is able to reduce the production of excessive ROS, maintaining the oxidation-reduction environment and the stability of mitochondrial membrane potential^13,14,15,16^. Interestingly, the formation of autophagosomes (mitophagosomes) can also occur at ERMCs and it has been shown that altering the structure of ERMCs can reduce the number of autophagosomes produced.^3,14,17^ Thus, it is likely that the establishment of ER-mitochondrial connections and the initiation of mitophagy are two inter-related processes.

Previous studies have shown that VAP27-1 localize to the ER-PM contact sites (EPCS) and the entire ER membrane^18,19,20,21,22,23^. VAP27-1 interacts with proteins involved in cytoskeleton interaction (e.g NET3C), membrane trafficking (e.g AtEH1/Pan, Clathrin), lipid transport (e.g ORP3a) and lipid-droplet biogenesis (e.g SEIPIN), regulating multiple subcellular activities. Here, we identify a novel mitochondrial membrane protein TraB1a, which also interacts with VAP27-1 to regulate ER-mitochondrial tethering. In parallel, TraB1 also interacts with ATG8 to regulate mitochondrial degradation functioning as putative mitophagy receptors. From a combination of experimental approaches, our data indicate that TraB1-mediated EMCS participates in mitophagy, which in turn maintains healthy mitochondrial function, and energy homeostasis. These findings not only advance our knowledge of organelle interactions and mitophagy in general, but also bridge the gap in our understanding of how EMCSs and mitophagy are regulated *in plantae*.

## Results

### TraB1 localizes to the mitochondria outer membranes and interacts with VAP27 at ER-Mitochondria contact sites

In eukaryotic cells, the endoplasmic reticulum (ER) contacts with most membranes^2,7,21,24,25,26,27,28^. Similarly, the ER is also connected to the mitochondria through ER-Mitochondria Contact Sites (EMCSs), which are essential in lipid metabolism, autophagy, mitochondrial function and morphogenesis^3,4,5,11,29^. Across eukaryotes, the ER-integral membrane protein VAP27 (also known as VAP/Scs2) serves as a tethering factor between the ER and various organelles^18,19,30,31^. We therefore investigated whether VAP27 may mediate contact sites between the ER and mitochondria in plants. In Arabidopsis lines co-expressing VAP27-1-GFP and the mitochondrial matrix marker, Mito-mCherry, we observed that some mitochondria partially co-localise with VAP27-1 on the ER surface (Figure S1a, arrow); such a phenomenon was not observed in control cells expressing the ER lumen marker, GFP-HDEL (Figure S1b), suggesting VAP27-1 may play an important role in mediating ER-mitochondria tethering in plants.

To screen for the tethering proteins through which VAP27 mediates EMCSs, TAP (Tandem affinity purification) tagging was performed using transgenic Arabidopsis cell cultures. After affinity purification followed by a MS proteomic screen, we identified TraB1a (Figure 1a), a mitochondrial protein with unknown function, as an interacting protein of VAP27-1. There are two closely related TraB1 proteins encoded by the Arabidopsis genome, TraB1a and TraB1b, both of which co-localize with VAP27-1-RFP at numerous donut-shaped structures when co-expressed (Figure 1a-b). These structures are likely to be a hybrid membrane consisting of the mitochondrial membrane and the ER membrane because both the GFP-TraB1a/TraB1b and VAP27-1-YFP signal co-localise at circular structures surrounding Mito-mCherry (Figure 1d). This phenomenon is likely caused by a direct interaction between TraB and VAP27-1, as it is absent in cells only expressing VAP27-1-RFP (Figure 1e).

**Figure 1.**
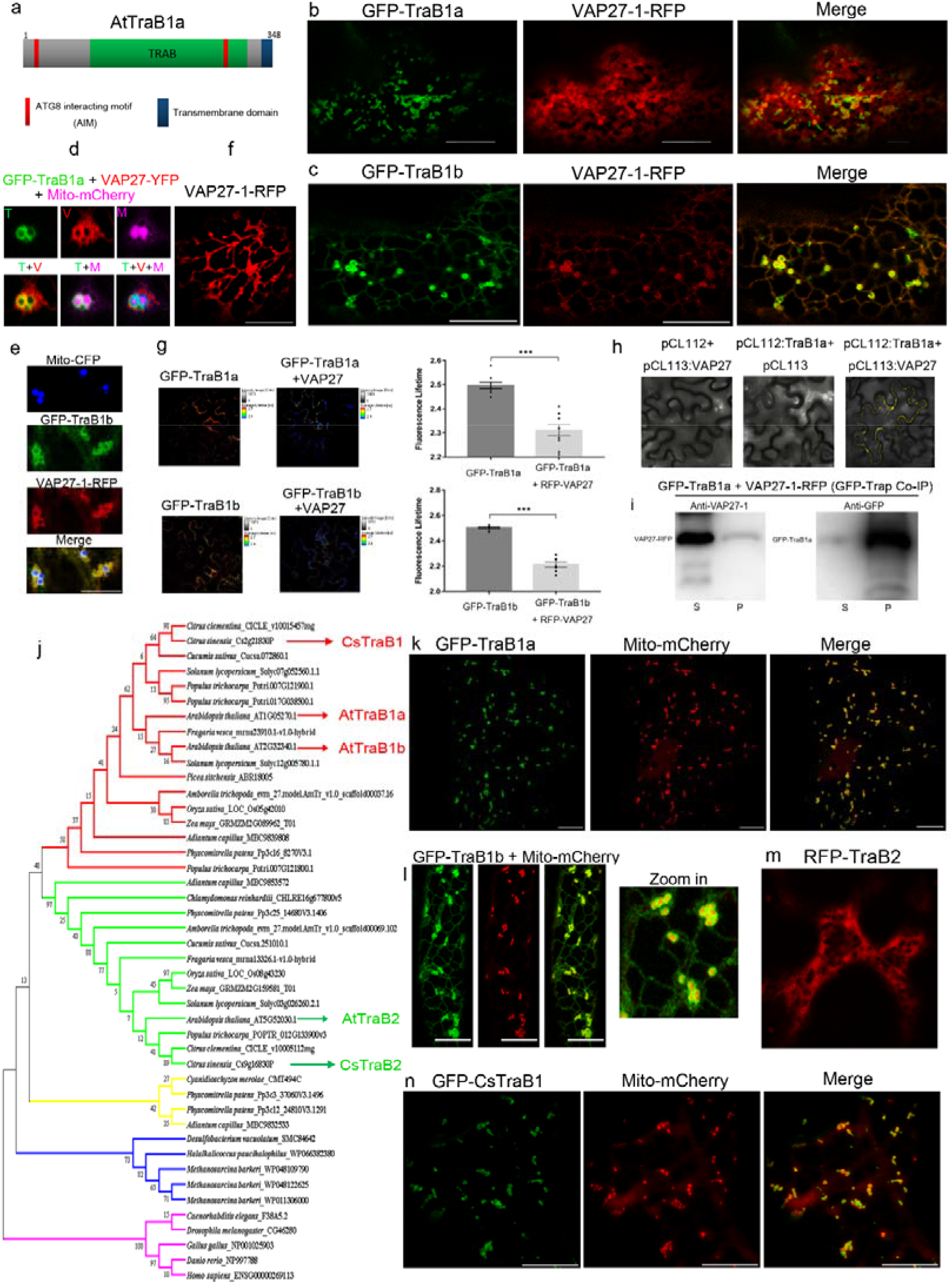
Phylogenetically conserved TraB1 proteins localise to mitochondria and interact with VAP27. **a**. Schematic illustration of TraB1a protein in Arabidopsis; it contains two ATG8 interacting motifs (AIM), one TRAB homology domain and a C-terminal transmembrane domain. **b**. GFP-TraB1a and VAP27-1-RFP co-localize at ER-derived donut-shaped membrane structures in *N. benthamiana* leaf epidermal cells. **c**. GFP-TraB1b co-localizes with VAP27-1-RFP at the ER network and donut-shaped membrane structures. **d**. GFP-TraB1a (green), VAP27-1-YFP (Red) and Mito-mCherry (magenta) co-expressed in *N. benthamiana* leaf epidermal cells, demonstrating that the ER derived donut-shaped structures superimpose with the mitochondrial outer membrane. **e**. GFP-TraB1b (green), VAP27-1-YFP (Red) and Mito-CFP (blue) co-expressed in *N. benthamiana* leaf epidermal cells, the ER derived donut-shaped structures superimpose with the mitochondrial outer membrane. **f**. Control cells expressing VAP27-1-RFP alone. **g**. FRET-FLIM further proved the interactions between TraB1a/VAP27-1 and TraB1b/VAP27-1. The fluorescence life-time of GFP-TraB1a (control) and GFP-TraB1b changes significantly (life-time reduces from 2.50 ± 0.01 to 2.31 ± 0.02 ns; and from 2.50 ± 0.01 to 2.21 ± 0.02 ns, respectively) in the presence of RFP-VAP27-1. **h**. Split-YFP based BiFC study showed nYFP-VAP27-1 and cYFP-TraB1 producing signals when co-expressed in *N. benthamiana*. **i**. The interaction between VAP27-1 and TraB1 is confirmed by a co-precipitation assay (GFP-Trap) using GFP-TraB1a and VAP27-1-RFP expressing plants. The majority of the GFP-TraB1a protein is trapped in the pellet fraction (left), where VAP27-1-RFP is also pulled-down in the presence of GFP-TraB1a (right). **j**. Phylogenetic tree of TraB homologues showing that Arabidopsis TraB1a and TraB1b originate from the same clade, whereas TraB2 belongs to a separate clade that is more evolutionarily similar to their mammalian homologues. **k**. In Arabidopsis, GFP-TraB1a localizes to the outer membrane of mitochondria that is labelled by the mitochondrial matrix marker, Mito-mCherry. **l**. GFP-TraB1b localizes to the ER network as well as the outer mitochondrial membrane. **m**. RFP-TraB2 mostly localizes to the cytoplasm. **n**. In citrus, only one isoform of TraB1 exists, which also localizes to the outer mitochondrial membrane in *N. benthamiana* leaf epidermal cells. N ≥ 10 for every FRET-FLIM analysis, error bars are SEM, *** P < 0.001 in Student’s t tests (Scale bar = 10 μm).

FRET-FLIM was used to further study these interactions. The fluorescence lifetimes of GFP-TraB1a and GFP-TraB1b are reduced significantly in the presence of RFP-VAP27-1, indicating physical interactions (Figure 1f-g). Furthermore these *in vivo* interaction data were further confirmed using co-immunoprecipitation assays, and also BiFC. Taken together we conclude that TraB1 and VAP27-1 interact (Figure 1h-i) and that this interaction may serve to mediate the formation of EMCS.

### Phylogenetic study of TraB proteins and their function in regulating mitochondrial activity

TraB1 proteins contain a C-terminal transmembrane domain, and a TraB-homology domain which currently has unknown function in eukaryotes (Figure 1a). In Arabidopsis, TraB1a and TraB1b are highly conserved, whilst TraB2 is more divergent (Figure 1j). GFP-TraB1a localises to ring-like structures that appear to be the outer membrane of mitochondria (OMM, Figure 1k), while the subcellular localisation of TraB1b, the closest homologue of TraB1 (sharing 73.4% peptide similarity), is found both at the ER network and the mitochondrial outer membranes (Figure 1i). These results are consistent with previous observations from a proteomic analysis^32,33^, where it was found that endogenous TraB1a is enriched in mitochondria in both Arabidopsis, and citrus fruits. Plant TraB2 isoforms (which are mainly localized to the cytoplasm; Figure 1m) are more closely related to the mammalian and bacterial TraB proteins, suggesting that this is likely to be an ancient form appearing earlier in evolution (Figure 1j). The mitochondrial localisation of TraB1 is likely to be conserved in many higher plants, as the citrus homologue of TraB1, CsTraB1 also localised to the OMM (Figure 1n).

TraB1a and TraB1b is expressed in most plant tissues with the strongest expression found in roots and flowers (Figure S2a-b). Under the control of their own promoters, expression of both GFP-TraB1a and GFP-TraB1b is further confirmed in vegetative tissues (Figure S2c-e). As they exhibit similar expression profiles and sequence homology, further studies were performed mainly using TraB1a as the example.

### TraB1 proteins are important in mitochondrial movement, morphology and energy metabolism

Endogenous TraB1a is found localized to the outer mitochondria membrane and stays closely associated with VAP27 in root cells (Figure 2a-b). This was demonstrated by immunofluorescence using an antibody that is specific to TraB1a (Figure S2f). These results confirm the live cell imaging data, suggesting that the endogenous TraB1 is also likely to facilitate the ER-mitochondria association. In the presence of over-expressed GFP-TraB1a or VAP27-1-RFP, the percentage of stationary mitochondria (that stay immobile for more than 6s, Figure 2c) increased significantly, likely due to an increment of ER-mitochondria tethering (Figure 2d-e). Enhanced ER-mitochondrial association at the ultrastructural level is observed in cells over-expressing GFP-TraB1a (Figure 2f-h). Taken together, we have identified a novel EMCS complex comprising the ER-localised VAP27 and the OMM-localised TraB1.

**Figure 2.**
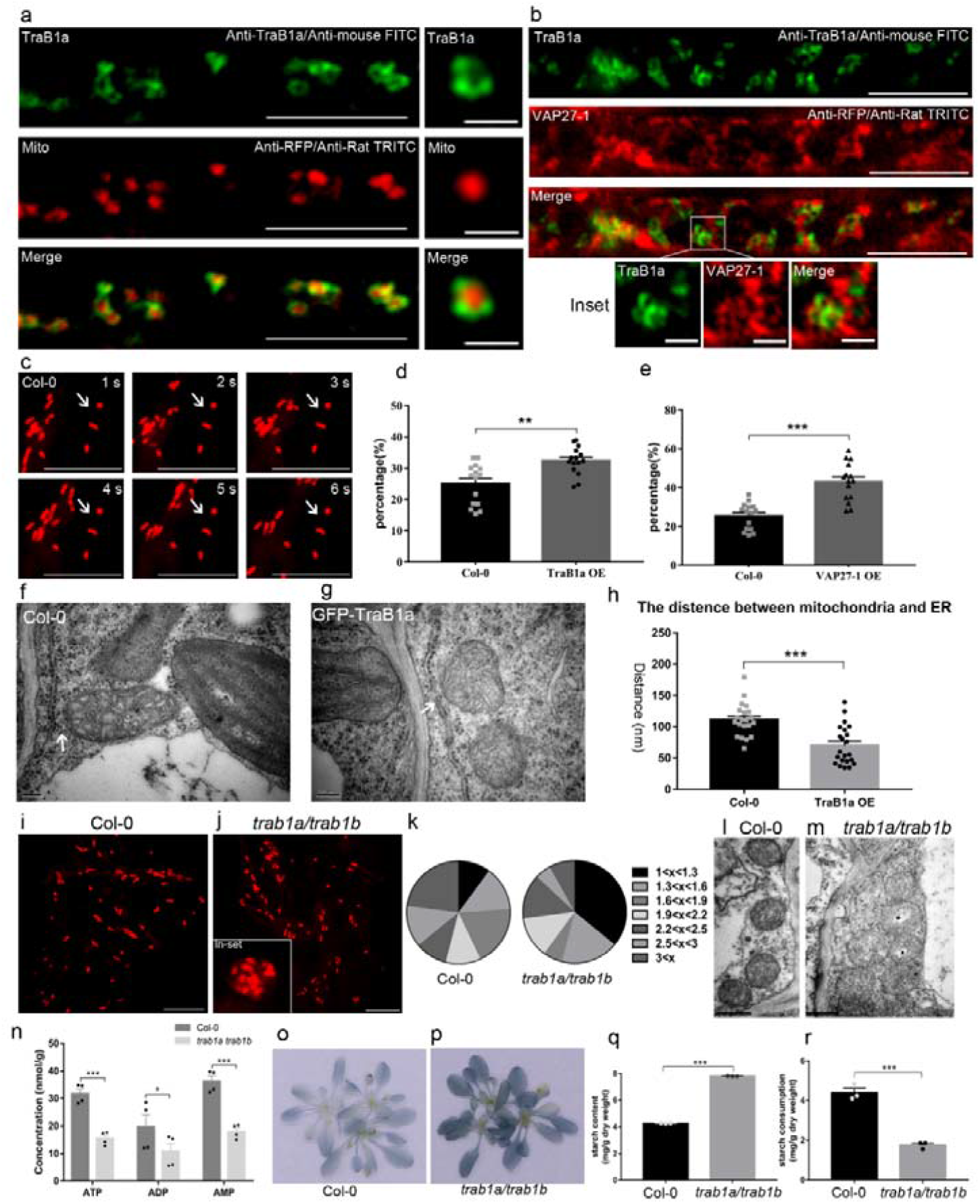
TraB1 proteins are essential for normal mitochondrial movement, morphology and energy metabolism. **a-b**. Arabidopsis expressing Mito-mCherry and VAP27-RFP was immuno-labelled with a TraB1a antibody; the endogenous protein localized at the outer mitochondrial membrane, and in contact with VAP27-labelled ER membrane, respectively. **c**. An example of stationary mitochondria (arrow) that stay immobile during the six second time course. **d-e**. The percentage of stationary mitochondria (immobile for at least six seconds) increased significantly in the presence of GFP-TraB1a or VAP27-1. **f-h**. TEM study of wild type Arabidopsis (f) and stable Arabidopsis transformed with GFP-TraB1 (g), the distance between ER and mitochondria is significantly reduced compared to wild type plants (h). **i-k**. In the *trab1a/trab1b* T-DNA mutant Arabidopsis line stably expressing Mito-mCherry, the morphology of mitochondria is significantly altered, as indicated by the increased population of circular mitochondria (aspec ratio at 1-1.3, k). Mitochondrial aggregations (j, inset) were also frequently observed. **l-m**. In the *trab1a/trab1b* Arabidopsis, mitochondrial structures are also disrupted at the ultrastructural level (m); their internal membrane cisternae are more fragmented and the matrix becomes more electron-transparent compared with that in the wild type (l). **n**. The levels of ATP, ADP and AMP are all significantly reduced in the *trab1a/trab1b* mutant, suggesting a disruption in energy metabolism and mitochondrial function. **o-q**. The starch levels in Col-0, *trab1a/trab1b* T-DNA plants (4-5 weeks old) were analysed by Lugol staining and the *trab1a/trab1b* loss-of-function mutant accumulated high levels of starch by the end of the light cycle (p). Such results were confirmed by spectrophotometry (q). **r**. Under dark conditions, starch consumption is reduced in the *trab1a/trab1b* mutant, suggesting a reduction in mitochondrial respiration. N ≥ 15 for mitochondria movement analysis; N ≥ 20 for TEM analysis; N = 9 for starch quantification; N ≥ 100 for mitochondria morphology analysis; N = 8for ATP measurement, error bars are SEM, * 0.01 < P < 0.05, *** P < 0.001 in Student’s t tests (Scale bar = 10 μm for light microscopy; Scale bar = 500 nm for TEM).

When TraB1a is over-expressed in *N. benthamiana* leaf epidermal cells, the mitochondria became enlarged and swollen (Figure S3a), and the morphology of mitochondria changes from a tubular/oval shape to a spherical shape (Figure S3b). Such phenotypes are commonly observed when the functions of mitochondria are inhibited^34,35,36^. At the ultrastructural level, these mitochondria appear to be aggregated and more electron transparent, suggesting mitochondrial function is likely impaired (Figure S3c-d). TMRM stains the mitochondria according to their electron potential, which can be used as an indicator for mitochondrial activity. In cells treated with antimycin (a disruptor of ATP synthesis), a significant reduction of TMRM fluorescence is seen (Figure S3e). Interestingly, the TMRM signal is weaker in Arabidopsis stably expressing GFP-TraB1a (Figure S3f) in contrast to control plants expressing Mito-YFP (Figure S3g), indicating that the expression of GFP-TraB1a results in decreased mitochondrial membrane potential and activity (Figure S3h-i).

### The function of mitochondria is affected in the *trab1a/b* mutant, indicated by a change in morphology, reduction of ATP and starch accumulation

Considering the high possibility of functional redundancy, the T-DNA double mutant of *trab1a/trab1b* was generated to further study the function of TraB1 (Figure S4a-b). The structure and morphology of mitochondria were analysed using light and electron microscopy. In the *trab1a/trab1b* T-DNA mutants, a large proportion of the mitochondria show a spherical shape and become aggregated which is in contrast to that observed in the wild type (Figure 2i-k). At the ultrastructural level, the structure of the mitochondria is disrupted in the *trab1a/b* mutants (Figure 2l-m). Moreover, the total ATP level and ATP/ADP ratio is reduced significantly in the double mutant (Figure 2n), indicating an aberrant energy metabolism.

In plants, starch is an energy reserve and its level in plants is closely related to energy homeostasis^37,38^. In agreement with the energy metabolism defect, we found strong starch accumulation in the *trab1a/trab1b* mutant (Figure 2o-q). In order to eliminate the influence from photosynthesis, and observe starch consumption directly related to mitochondrial activity, we further analysed the change in starch levels under dark conditions. We found a decrease in the reduction of the starch level in the *trab1a/trab1b* mutants compared to the wild type (Figure 2r), suggesting starch consumption is inhibited. Meanwhile, another independent *trab1a/b* double mutant was also created using CRISPR/Cas9 (Figure S5a) and a similar starch phenotype was observed (Figure S5c-d). In summary, these observations are consistent with the functions of TraB1 in regulating mitochondrial activity and ER-mitochondria tethering, since mitochondrial function and integrity are regulated by EMCSs^39,40,41,42^.

### TraB1 interacts with ATG8 through the conserved AIM motif

As the molecular functions of TraB1 are yet uncharacterised, we performed further sequence analysis and we found two ATG8-interacting motifs (AIM) in both proteins, suggesting it may interact with ATG8 and participate in the autophagy pathway^43,44^. Therefore, stable Arabidopsis lines expressing GFP-ATG8a and RFP-TraB1a were generated. We found that ATG8a signal is recruited to TraB1 labelled mitochondria under normal growth conditions (Figure 3a); such co-localisation is rarely found in cells expressing CFP-ATG8a and Mito-mCherry (Figure 3b).

**Figure 3.**
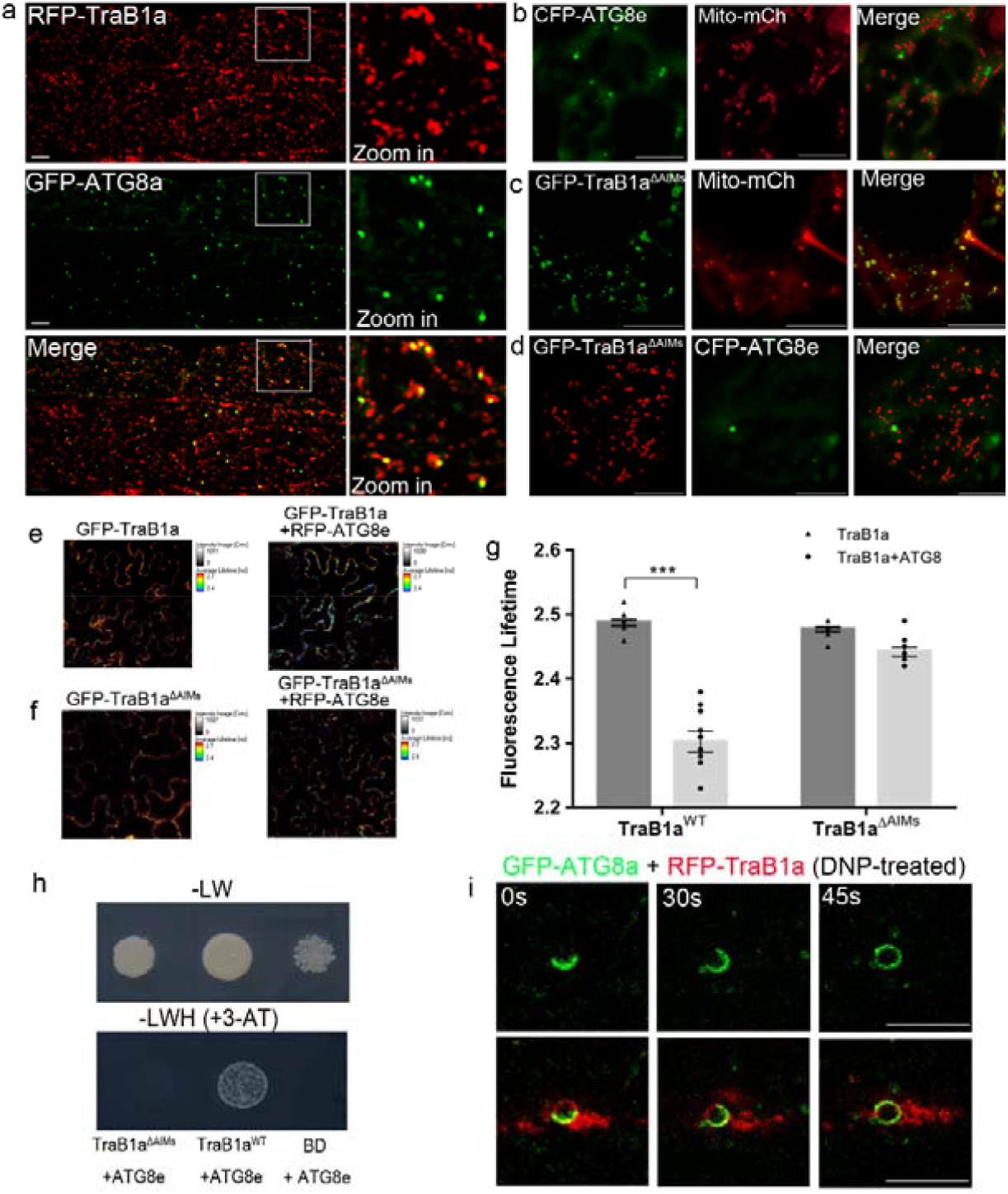
TraB1 interacts with ATG8 through the ATG8 interacting motif (AIM). **a**. In Arabidopsis stably transformed with RFP-TraB1a and GFP-ATG8e under non-stressed conditions, GFP-ATG8e positive structures are recruited to the mitochondria that are labelled with RFP-TraB1 in hypocotyl epidermal cells. **b**. Such ATG8e-mitochodrial co-localization is rarely seen in cells expressing a mitochondrial maker, Mito-mCherry. **c-d**. *N. benthamiana* leaf epidermal cells expressing GFP-TraB1ΔAIMs (TraB1a without the ATG8 interacting motif) with a mitochondrial marker (c) and an autophagosome marker (d). GFP-TraB1ΔAIMs still localizes to the mitochondrial outer membrane but did not co-localise with ATG8. **e-g**. FRET-FLIM further proved that the interactions between TraB1a and ATG8 is reliant on the AIM motifs. The fluorescence life-time of GFP-TraB1a (control) changes significantly (life-time reduces from 2.49 ± 0.01 to 2.30 ± 0.02 ns) in the presence of RFP-ATG8e, while the fluorescence life-time of GFP-TraB1ΔAIMs (control) only dropped by 0.04 ns in the presence of RFP-ATG8e; such a difference is too little to indicate an interaction. **h**. Using a one-on-one yeast-2-hybrid assay, ATG8e was confirmed to interact with full length TraB1a but not with the TraB1ΔAIMs mutant. i. Following DNP-induced mitochondrial depolarization in Arabidopsis root cells expressing GFP-ATG8e and RFP-TraB1a, the ATG8e labelled autophagosome membrane was recruited to a swollen mitochondrion that is labelled by TraB1a during a course of 45 seconds. N ≥ 10 for every FRET-FLIM analysis, error bars are SEM, * 0.01 < P < 0.05, *** P < 0.001 in Student’s t tests (Scale bar = 10 μm).

Next, we mutated the putative AIM motifs of TraB1a (denoted as TraB1aΔAIMs, Figure S4c). GFP-TraB1aΔAIMs localizes to the mitochondria like the full-length protein (Figure 3c). However, its co-localization with ATG8 was found to be significantly reduced (Figure 3d). FRET-FLIM was used to study the interaction of TraB1aΔAIMs with ATG8 (Figure 3e-g). We observed that the fluorescence lifetimes of GFP-TraB1a are reduced significantly in the presence of RFP-ATG8e, indicating physical interactions, whereas the GFP lifetimes exhibit little reduction when GFP-TraB1aΔAIMs is expressed with RFP-ATG8e using the same conditions (Figure 3g), indicating that GFP-TraB1a ΔAIMs does not interact with RFP-ATG8e. Similarly, co-localization and interaction between TraB1b and ATG8 was confirmed *in vivo* (Figure S6a-c). Using one-on-one yeast-two-hybrid assays, the yeast strains only grew on selective medium when co-transformed with non-mutated TraB1 and ATG8, but were not able to grow when co-transformed with TraB1aΔAIMs and ATG8 (Figure 3h). These data indicate that the AIM motif is essential for the interaction between TraB1a and ATG8e.

It is known that DNP promotes mitochondrial depolarisation and mitophagy^45,46,47^. After DNP treatment, time lapse confocal microscopy revealed the dynamic recruitment of ATG8 labelled autophagosome membranes to the swollen mitochondria in the presence of TraB1a (Figure 2i) which is consistent with a role for TraB1 in its interaction with ATG8.

### The turnover of TraB1 is dependent on the autophagy pathway

Concanamycin A (ConcA) is a V-ATPase inhibitor that inhibits lytic vacuole activities and induces autophagosome accumulation (Figure 4a-b)^48^ and we have used it to determine the autophagy flux of GFP-TraB1a under different conditions. In lines co-expressing RFP-TraB1 and GFP-ATG8a, large numbers of the TraB1 labelled puncta co-localize with ATG8 (Figure 4c-d) after Conc A treatment, suggesting that TraB1 is targeted to the vacuole with the autophagosomes. The number of vacuole-accumulated autophagosomes increases significantly in the presence of GFP-TraB1 (Figure 4e). Similarly, in cells expressing GFP-TraB1a and Mito-mCherry, both proteins co-localize in the vacuole after Conc A treatment (Figure 4f-i), and the number of vacuole-accumulated mitochondria increased significantly (Figure 4j) in contrast to the control cells only expressing Mito-mCherry (Figure 4f-g). These data suggest a possible function for TraB1 in promoting autophagy/mitophagy. Furthermore, in transgenic Arabidopsis expressing both GFP-TraB1b and Mito-mCherry, strong co-localisation and vacuole accumulation of these proteins was found after Conc A treatment (Figure S6d-e).

**Figure 4.**
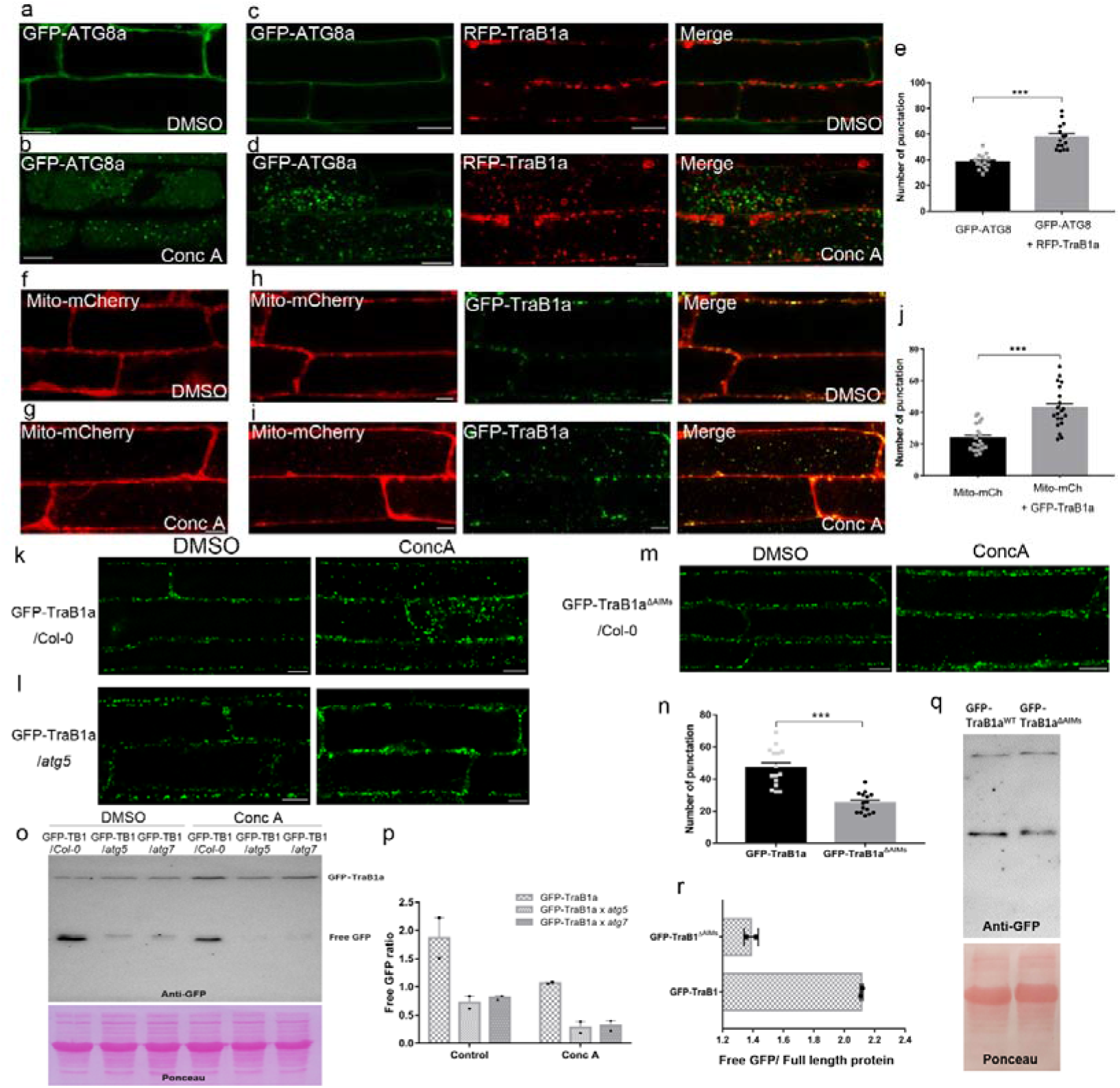
TraB1a over-expression promotes autophagy flux and the degradation of mitochondria through the autophagy machinery. **a-d**. Transgenic Arabidopsis plants expressing GFP-ATG8a or GFP-ATG8a+RFP-TraB1a grown in MS growth media without stress. After Conc A treatment, GFP-ATG8a labelled autophagosome structures were identified inside the vacuole (b); autophagosomes that contain mitochondrial signal (labelled by RFP-TraB1a) are also abundant in cells expressing GFP-ATG8a and RFP-TraB1a (d). In contrast, little vacuole signal was seen in cells treated with DMSO as the control (a, c). **e**. The number of autophagosomes in the vacuole after conc A treatment was analysed statistically; more autophagosomes were found in cells over-expressing TraB1a. **f-i**. Transgenic Arabidopsis plants expressing Mito-mCherry or Mito-mCherry + GFP-TraB1a grown in MS growth media without stress. After Conc A treatment, numerous mitochondria (either labelled with Mito-mCherry or co-labelled with Mito-mCherry + GFP-TraB1a) accumulated in the vacuole (g,i). In contrast, little vacuole signal was seen in cells treated with DMSO as the control (f, h). **j**. The number of vacuole-accumulated mitochondria increased in cells over-expressing TraB1a after Conc A treatment. **k-l**. GFP-TraB1a localization in wild type and an autophagy deficient mutant (*atg5*) under non-stressed conditions upon Conc A treatment. No punctate structures are found to accumulate inside the vacuole of cells in the *atg5* mutant, indicating that the transport of TraB1-labelled mitochondria into the vacuole relies on the core autophagy machinery. **m-n**. Stable Arabidopsis expressing GFP-TraB1ΔAIMs were treated with Conc A (m), the vacuole-accumulation of GFP positive signal is significantly reduced in comparison with Arabidopsis expressing non-mutated GFP-TraB1a (n). **o-p**. Western blot analysis of GFP-TraB1 (GFP-TB1) levels in *Col-0, atg5* and *atg7* Arabidopsis seedlings with and without Conc A treatment (o), the result shows a reduced level of degradation of GFP-TraB1 in both of the autophagy defective mutants, as indicated by the reduction in free GFP levels and ratio of free GFP/Full-length protein (p). **q-r**. Western blot analysis of the protein levels of GFP-TraB1 and GFP-TraB1ΔAIMs in Arabidopsis seedlings under non-stressed condition (q), the result shows that the degradation of GFP-TraB1ΔAIMs is reduced, as indicated by the reduction in free GFP level and ratio of free GFP/Full-length protein (r). N ≥ 15 for image quantification; N = 2 for every western blot analysis, error bars are SEM, *** P < 0.001 in Student’s t tests (Scale bar = 10 μm; Scale bar = 15 mm for root analysis).

The internalization of GFP-TraB1a to the vacuole is dependent on autophagy, as no vacuole accumulation of TraB1a is found after Conc A treatment of the same GFP-TraB1a transgenic line crossed into either the *atg5* and *atg7* autophagy deficient mutants (Figure 4k-l). In parallel, we generated Arabidopsis transgenic lines expressing the ATG8 binding-defective GFP-TraB1aΔAIMs mutant construct, in which little GFP-TraB1aΔAIMs vacuole accumulation is observed following Conc A treatment (Figure 5m-n). Therefore, these results suggest that the vacuole internalization of TraB1 is dependent on both the interaction with ATG8 and a functional autophagy pathway.

**Figure 5.**
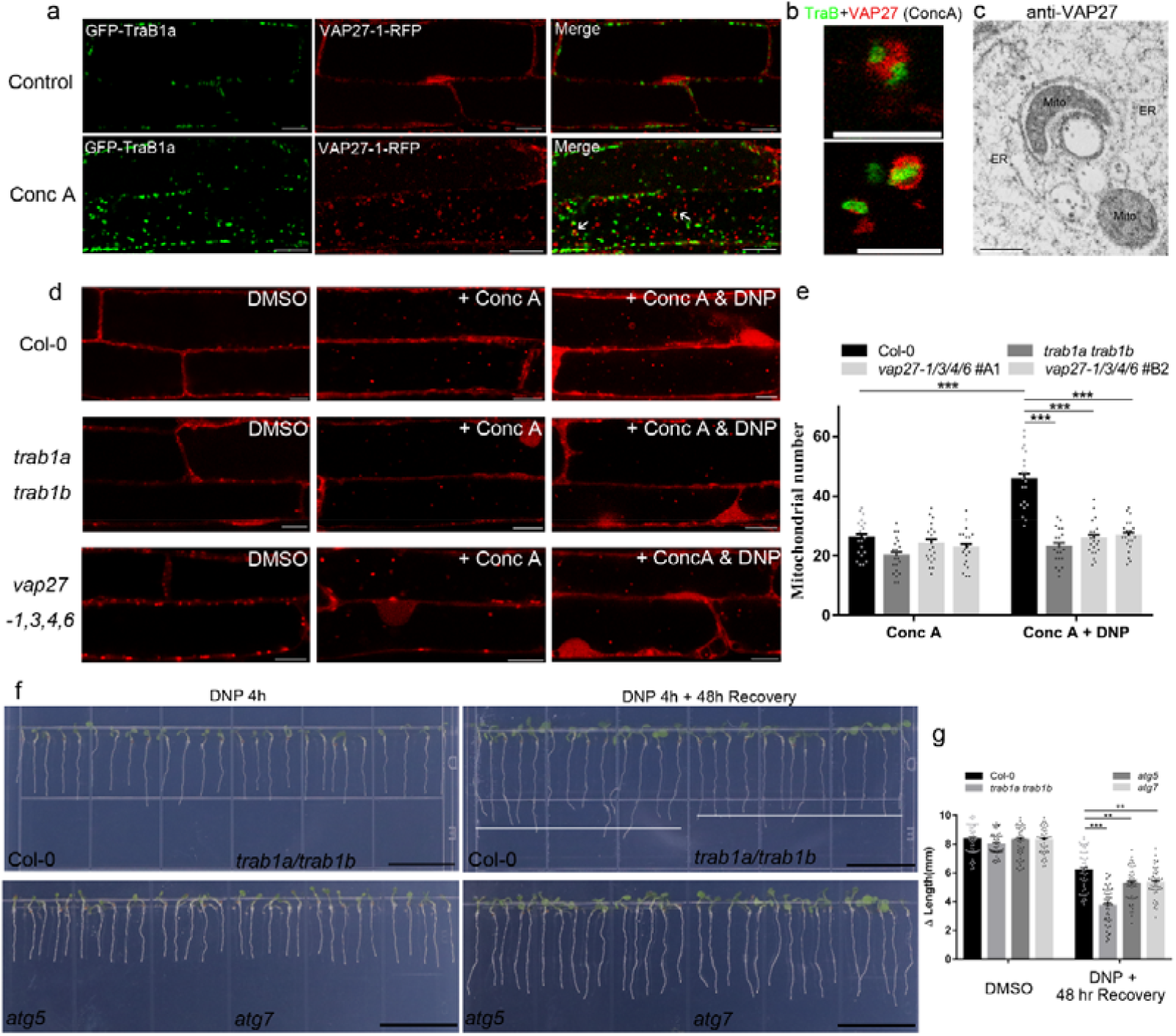
Mitochondrial stress induced mitophagy is inhibited in the *trab1a/trab1b* mutants. **a**. Transgenic Arabidopsis expressing GFP-TraB1 and VAP27-1-RFP were treated with Conc A. The majority of TraB1-labelled mitochondria are found in contact with VAP27-labelled ER membrane inside the vacuole, suggesting that these two structures are internalised together. **b**. High magnification images of **a. c**. Anti-VAP27 immunogold labelling of transgenic Arabidopsis Hypocotyl cells expressing VAP27 under non-stressed condition. **d-e**. Arabidopsis plants (Col-0; *trab1a/trab1b; vap27-1,3,4,6*) stably expressing Mito-mCherry were treated with DNP and Conc A to study the activity of mitophagy. Under non-stressed conditions, the mitochondrial turnover rate is similar in all plants analyzed. However, the number of vacuole-accumulated mitochondria is significantly reduced in both the *trab1a/trab1b* and the *vap27-1,3,4,6* (two independent lines, A1 and B2) mutants treated with DNP, suggesting that an interruption of ER-Mitochondria tethering affects mitochondrial turnover following mitochondrial stress/mitochondrial damage. **f-g**. The recovery of Arabidopsis seedlings after 4 hours of DNP treatment was studied. The root growth of the trab1a/b mutant plants is significantly reduced compared to that in the wild type, suggesting that the DNP induced mitochondrial dysfunction has more impact on plant development when the function of TraB is impaired. N ≥ 20 for confocal quantification; N ≥ 90 for root analysis; error bars are SEM, *** P < 0.001 in Student’s t tests (Scale bar = 10 μm for confocal; Scale bar = 200 nm for TEM).

To further confirm the autophagic degradation of TraB1, protein levels of GFP-TraB1 in autophagy deficient backgrounds was monitored using western blotting^48^. The levels of full-length protein and their degradation products (free GFP cleaved from the fusion protein) were analysed. As expected, the level of full-length GFP-TraB1a protein was found to be higher in *atg5, atg7* mutants under normal growth conditions, whilst the level of free GFP resulting from protein degradation was much lower compared to the Col-0, indicating impairment of TraB1 degradation in the autophagy mutants (Figure 5o-p). When treated with Conc A, the protein levels of GFP-TraB1 accumulated to a higher level in the wild type, indicating that blocking autophagic degradation prevents TraB1 degradation. Meanwhile, ConcA treatment of both *atg5* and *atg7* mutants did not result in a pronounced accumulation of GFP-TraB1a levels (Figure 5o-p). Transgenic plants expressing GFP-TraB1aΔ AIMs were also analysed and we found that the intensity ratio of free GFP/full-length protein is reduced significantly in GFP-TraB1ΔAIMs plants (Figure 4q-r), indicating the impaired degradation of TraB1 when its interaction with ATG8 is inhibited.

Interestingly, the impairment of autophagic degradation of GFP-TraB1 also resulted in developmental defects. Both *atg5* and *atg7* plants expressing GFP-TraB1 are smaller than Col-0 lines expressing GFP-TraB1 (Figure S7). Similarly, Arabidopsis plants expressing GFP-TraB1ΔAIMs are also dwarfed (Figure S8), indicating that the autophagy-dependent protein turnover of TraB1 is essential for normal plant development, and over-accumulation may lead to mitochondrial dysfunction (Fig.S3a-d). Taken together, in addition to the function as an ER-Mitochondria tethering factor by its interaction with VAP27-1, TraB1 proteins also interact with ATG8 and are degraded via the autophagy pathway. However, whether TraB1 is part of the machinery for mitophagy, or simply acts as a cargo for autophagic degradation is the next question to be determined.

### Stress induced mitophagy is affected in the *trab1a/b* mutant, suggesting that TraB1 is likely to be a mitophagy receptor

It is known in yeast and mammalian systems that the ER-Mitochondrial interfaces are essential for autophagosome biogenesis and mitophagy^17,49,50^, so it is also reasonable to propose that TraB1-VAP27 mediated ER-Mitochondrial tethering and TraB1-ATG8 regulated mitophagy are two closely related processes. Transgenic Arabidopsis expressing GFP-TraB1 and VAP27-RFP were treated with Conc A. The majority of the TraB1 labelled mitochondria are found in close contact with VAP27 labelled ER membrane after internalization to the vacuole, suggesting that ER-Mitochondria stay in contact during mitophagy (Figure 5a-b). With the help of immunogold labelling, anti-VAP27 gold particles were found to be enriched at the ER-Mitochondrial interface of mitophagosome-like structures (Figure 5c).

To investigate whether TraB1 and VAP27 are important for regulating mitophagy, we examined mitophagy activity in their mutant plants. Mito-mCherry was transformed into wild type Arabidopsis and the *trab1a/b* double mutant, and the level of mitophagy flux (indicated by the number of internalized mitochondria) was measured after ConcA treatment. Under normal growth conditions, the number of vacuole-internalized mitochondria was found to be similar in both Col-0 and *trab1a/b* but a significant reduction was observed in the *trab1a/b* mutant when these plants were subjected to DNP-induced mitochondrial depolarization (Figure 5d). This result indicates that TraB is needed for stress-induced mitophagy above the normal basal levels. In addition, two independent *vap27-1/3/4/6* quadruple mutants were generated (Figure S9a) using CRISPR/Cas9 with the intention of disrupting the VAP27-TraB1 interaction at EMCSs, as these ER localized VAP27s exhibit partial co-localization with TraB1a (Figure S9b-d). Both mutants exhibit defects in stress-induced mitophagy similar to the *trab1a/b* mutant (Figure 5d-e).

We hypothesise that heightened levels of impaired mitochondria (TMRM negative) resulting from DNP-induced depolarisation (Figure S10a) requires mitophagy to maintain mitochondrial homeostasis, in which TraB1-VAP27 plays an important role. Significant accumulation of depolarized mitochondria is found in the loss-of-function mutants of *trab1a/b* and *vap27-1/3/4/6* (Figure S10b-d). In conclusion our data demonstrate that TraB1: 1. is OMM-localized; 2. interacts with ATG8; 3. is transported to the vacuole using the autophagy machinery; 4. over-expression enhances mitophagy and furthermore, 5. in TraB1 knock out mutants, DNP induced mitophagy is partially impaired. These data favour the hypothesis that TraB1 is a novel mitophagy receptor, as well as performing a function in ER-Mitochondria tethering.

Mitochondrial damage releases reactive oxygen species and other toxic compounds^16,51^, which induces cell stress. Therefore, unsuccessful removal of damaged/depolarised mitochondria may have negative impacts on plant growth^52,53^. To test this hypothesis, the Arabidopsis *trab1a/b* and *vap27-1/3/4/6* mutants were subjected to acute DNP treatments and their growth was analysed 2 days after recovery on DNP-free media. We observed that *trab1a/b* and *vap27-1/3/4/6* mutant plants exhibit relatively weak recovery compared to Col-0 plants after 2 days (Figure 5f, Figure S11), suggesting that both mutants are defective in the maintenance of mitochondrial homeostasis, which we attribute to defects in mitophagy. This observation is consistent with our previous observations that impairment of TraB-promoted mitophagy (through overexpression of GFP-TraB1ΔAIMs) results in growth defects (Figure S8), and demonstrates the importance of endogenous TraB in plant mitophagy. As the positive control, the autophagy defective mutants (*atg5* and *atg7*) were treated with DNP and studied in the same way; reduced recovery was also prominent (Figure 5f-g).

## Discussion

It is known that ER-Mitochondria interactions are essential for mitochondrial division and signal/substrate exchange between the two compartments. In animal cells, the protein composition of EMCSs has been extensively studied in the last decade, and proteins such as MFN, Miro and dynamin related proteins are known to have important functions in regulating mitochondrial function and morphogenesis^11,36,42,49,54,55^. In this study, we have identified the evolutionarily conserved TraB1 protein family as regulators of two putative interrelated pathways. Firstly, they interact with ER-localized VAP27 to regulate ER-Mitochondrial interactions, and secondly, they interact with ATG8 to regulate mitochondrial degradation working as putative mitophagy receptors. Our data indicates that TraB1 has important roles in mitochondrial homeostasis through regulation of mitophagy, which is essential for normal plant growth under stress conditions.

It is known in yeast and mammalian systems that the ER-Mitochondrial interfaces are essential for autophagosomes biogenesis^3,17,40,42,49,50,56^, so it is also reasonable to propose that TraB1-VAP27 mediated ER-Mitochondria tethering and TraB1-ATG8 regulated mitophagy are two closely related processes. In animal cells, the homologues of VAP27 (VAPA/B) interact with multiple autophagy proteins, including ULK1/ATG1 and WIPI2/ATG18, to modulate autophagosome biogenesis^26^. In plants, VAP27 also functions in regulating actin-dependent autophagosome biogenesis through recruitment of AtEH1/Pan1^21^. Therefore, it is likely that the core autophagy machinery can also be recruited to the VAP27-mediated membrane interfaces, the VAP27-TraB1-mediated EMCSs, and function in selective autophagy processes (Figure 6).

**Figure 6.**
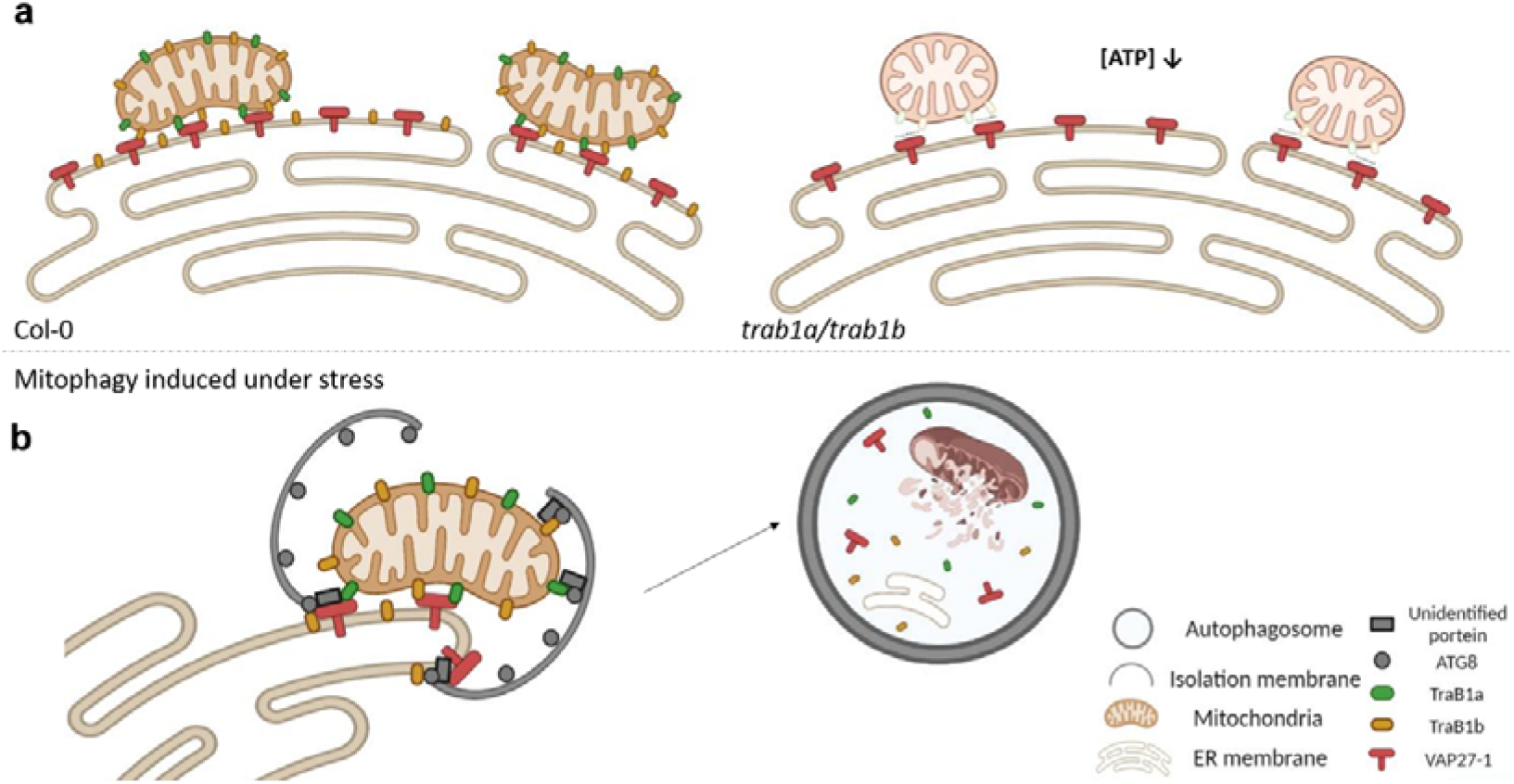
Proposed model for TraB1-VAP27 mediated Mitochondria-ER tethering and mitophagy. **a**. Under normal conditions, mitochondria are tethered to the ER through the TraB1-VAP27 complex at EMCSs; this link is essential for mitochondrial function (e.g. movement, morphology and energy metabolism). **b**. TraB1 also interacts with ATG8, a function that is essential for the mitochondrial stress-response. When the number of damaged mitochondria increases, TraB1 recruits ATG8 at the EMCS to regulate the formation of mitophagosomes. VAP27 may also recruit other unidentified autophagy proteins, such as ULK1/ATG1 and WIPI2/ATG18 to facilitate autophagosome maturation, as has been reported in mammalian cells.

In the *trab1a/b* mutant, mitochondrial structure and function are impaired, causing an overall reduction in ATP production and starch consumption (Figure 3n-r). These phenotypes are indicative of defective mitochondrial function and are likely linked to the function of TraB1 in mitophagy and ER-Mitochondria tethering, both of which are important in maintaining mitochondrial homeostasis and a healthy population of mitochondria through turnover of damaged mitochondria. In animal cells, it has been observed that damaged sections of mitochondria become dissected from the entire mitochondrial network, and are degraded through mitophagy. Unsuccessful removal of damaged mitochondria causes overall mitochondria dysfunction and cell stress/death^10,13,47,52,53,57,58^. Consistent with this, TraB1-loss of function results in the accumulation of damaged mitochondria under stress conditions caused by DNP treatment (Figure 5d, Figure S11).

In conclusion, we have characterised a novel EMCS complex mediated by the interaction of VAP27 with the OMM-integral protein, TraB1, which has important roles in mitochondrial function through regulating ER-Mitochondrial tethering and mitophagy (Figure 6a, b). Firstly, TraB1 interacts with ER-localized VAP27 to regulate ER-Mitochondrial interactions. Secondly, TraB1 interacts with ATG8 to regulate mitochondrial degradation working as putative mitophagy receptors, which is essential for normal plant growth in stressed conditions. We hypothesise that TraB1-mediated EMCSs are likely to regulate mitophagy, which in turn maintains healthy mitochondrial homeostasis and function. Whilst the mechanisms of EMCSs are already well known in animals and yeast, TraB1 is the first identified protein to have a dual function in ER-mitochondria tethering and mitophagy in eukaryotes, and these data aid in bridging the gap in our understanding of how EMCSs and mitophagy are regulated *in plantae*.

## Materials and methods

### Molecular cloning

The cDNA of TraB1a, TraB1b and TraB2 were cloned into pMDC43 or pK7WGR2 via gateway cloning (BP Clonase™ II, Invitrogen 11789020; LR Clonase™ II, Invitrogen 11791020), to facilitate their expression as an N-terminal GFP or RFP fusion proteins. ATG8e was cloned using Phanta Max Super-Fidelity DNA Polymerase (Vazyme, P505) into pK7WGR2 (N-terminal RFP fusion protein) and pK7WGC2 (N-terminal CFP fusion protein) via gateway cloning. To make the AIM motif mutation of TraB1a, two predicted AIM motifs (Figure S4c) were mutated (F21A, I24A and F279A, L282A) using their site-mutagenesis primers and a Mut Express II Fast Mutagenesis Kit-V2 (Vazyme, C214).

For the expression of TraB1a and TraB1b under their own promoters, *pTraB1a:*GFP-TraB1a and *pTraB1b*:GFP-TraB1b native expression constructs were generated. Approximately 1500bp of promoter region directly upstream of the TraB1a and TraB1b start codons was PCR amplified with 5’ PstI and 3’ BamHI restriction sites using their gene specific primers. The *pCAMV 35s*: promoters of pMDC43-TraB1a and pMDC43-TraB1b were excised by PstI/BamHI double restriction digest, and PstI/BamHI-digested *pTraB1a* and *pTraB1b* fragments were inserted respectively using T7 DNA Ligase (NEB, M0318S). For generation of recombinant TraB1a-6xHIS protein, full length TraB1a cDNA was PCR amplified with 5’ NdeI and 3’ NotI restriction sites using gene-specific primers, and subcloned into the NdeI/NotI restriction sites of pET24 using T7 DNA ligase (NEB, M0318S) to facilitate its expression as a C-terminal 6xHIS fusion protein in *E. coli*.

For yeast-two-hybrid constructs, full-length CDS of ATG8e was sub-cloned into pGADT7-GW plasmid using Gateway. TraB1a and TraB1aΔAIMs coding sequences without the transmembrane domains were cloned into pGBKT7 plasmid by Gateway LR reaction. All primers used in this study were listed in Supplementary Table S1.

### Transgenic Arabidopsis lines

Mito-CFP, Mito-YFP, Mito-mCherry, GFP-HDEL organelle markers have been described previously^59^. To generate the GFP-ATG8a and RFP-TraB1a Arabidopsis lines, the GFP-ATG8a Arabidopsis line^21,60^ was dipped with RFP-TraB1a via Agrobacterium-mediated transformation. Arabidopsis lines co-expressing GFP-TraB1a + Mito-mCherry, GFP-TraB1a + VAP27-1-RFP and GFP-TraB1b + Mito-mCherry were generated by crossing using stable transgenic plants of expressing single constructs. The GFP-TraB1a/*atg5*, GFP-TraB1/*atg7* were generated by crossing of stable transgenic GFP-TraB1a plants and the *atg5* and *atg7* mutants. Other Arabidopsis mutants generated in this study and are described in detail below. Please refer to Supplementary Table S2 and S3 for a complete list of all plasmid and transgenic Arabidopsis lines in this study.

### Arabidopsis Mutant analysis

The *trab1a-1* (GABI KAT_369F08) T-DNA mutant line was obtained from GABI Kat, and the *trab1b-1* (SALK_059433) T-DNA mutant line was obtained from NASC, and the *trab1a-2, trab1b-2*, and *vap27-1,3,4,6* CRISPR mutants were generated in this study. The *atg5* (SAIL_129_B07) and *atg7* (SAIL_11_H07) mutants have been described as previously^61^.

For T-DNA mutant lines, homozygosity was confirmed using the genotyping primers listed in Supplementary Table S1. Total RNA was extracted from seedlings (Qiagen) and cDNA was synthesised using Superscript III (Invitrogen). RT-PCR was used to confirm the absence of full-length transcripts in the T-DNA mutants using the primers listed in Supplementary Table S1. RT-PCR amplification of the housekeeping gene, *EF1⍰*, was performed as a control, and equal amounts of Col-0 and mutant cDNA was used.

For CRISPR/Cas9-mediated generation of *trab1a* and *trab1b* mutants, two gRNA spacer sequences on the + strand were chosen as Cas9 targets: for TraB1a; target 1 (5’-GCCGCCGCTAAACACAGAAT-3’) and target 2 (5’-GAAGACAGCTTGAAACAGTA-3’) were chosen. For TraB1b; target 1 (5’-GGAGAAGTTGGAGCTGCCTG-3’) and target 2 (5’-ATTGCAGATAAAGGAACTGA-3’) were chosen. For *vap27-1/vap27-3* double mutants, two gRNA spacer sequences on the + strand of each gene were chosen as Cas9 targets. For VAP27-1, target 1 (5’-AGGTCTACTTGCGAAGTTCT-3’) and target 2 (5’-ATACTGGAGTTGTTCTCCCG-3’) were chosen; for VAP27-3, target 1 (5’-CGATAATTATGTCGCCTTCA-3’ and target 2 (5’-ATCCAAAGAAGTACTGCGTT-3’) were chosen. Target specificities were evaluated with CAS-OFFINDER, using an algorithm for potential off-target sites of Cas9 RNA-guided endonucleases (Bae et al., 2014). The two respective TraB1a and TraB1b gRNA expression modules, were constructed via PCR using pCBC-DT1T2 as a template according to Xing et al.^62^ and Golden Gate assembled into the vector pHEE401E allowing egg cell-specific EC1.2en:EC1.1 promoter-controlled expression of 3× FLAG-NLS-zCas9-NLS^63^. The four VAP27-1/VAP27-3 gRNA expression modules were constructed via PCR using pCBC-DT1T2, pCBC-DT2T3 and pCBC-DT3T4 as a template according to Xing et al.^62^ and assembled via HiFi Gibson Assembly into pHEE401E. For *vap27-1,3,4,6* CRIPSR mutants, the two VAP27-4/VAP27-6 gRNA expression module was transformed into *vap27-1/vap27-3* CRISPR mutants generated above. The two gRNA expression module was constructed via Restriction-ligation cloning into pKSE401^63^, and the target (5’-TGGGACTCTTGCATCATTAG-3’) of VAP27-4, target (5’-GGACGAACACAGTATTTGCG-3’) of VAP27-6 were chosen respectively. Transformants were genotyped and sequenced using the primers listed in Supplementary Table S1.

### Generation of antibodies and immunostaining

Polyclonal antibodies were raised to full-length TraB1a-6xHIS. TraB1a-6xHIS was expressed in *E. coli* (Rosetta2; Novagen), and purified using Nickel-Agarose beads (Qiagen). Polyclonal antibodies were raised in mice as previously described^64^.

For immunostaining, 5–7-day old seedlings were fixed in 4 % PFA in PEM buffer (50 mM PIPES pH 6.9, 5 mM EGTA, 1 mM MgSO_4_) for 90 minutes. Fixed roots were washed (5x 10 min) with PEM buffer and cell walls were digested using 0.5 % cellulase, 0.05 % pectolyase, 2 % dricelase in PEM buffer + 0.4 M mannitol for 15 minutes. Roots were washed (3x 10 minutes in PEM) and were gently squashed on a poly-L-lysine-coated coverslip. Roots were washed (3x 10 minutes in PEM) and permeablised in PEM + 0.1% triton X100 for 15 minutes. Roots were washed (3x 10 minutes in PEM) and incubated in blocking buffer 2% BSA in PEM) for 1 hour. Samples were incubated in primary antibody at room temperature for one hour and then at 4°C overnight. Samples were then washed in Phosphate Buffered Saline solution (PBS; 6x 30 minutes) and were stained with secondary antibody at room temperature for one hour and then at 4°C overnight. Samples were washed in PBS (6x 30minutes) and were mounted in Vectashield and imaged using Airy scanning super-resolution confocal microscopy.

For co-staining of TraB1a with VAP27-1 and mitochondria, Arabidopsis stable lines expressing native VAP27-1-RFP or Mito-mCherry were stained with anti-TraB1a mouse primary antibody at a dilution of 1:500 and anti-RFP rat primary antibody (Chromotek), followed by secondary antibody incubation with FITC conjugated against mouse, and TRITC conjugated against rat (Jackson ImmunoResearch).

### Live cell Imaging and Image analysis

*Nicotiana benthamiana* plants were grown in a growth room or greenhouse with long-day conditions. Transient expression was performed by leaf infiltration according to Sparkes et al^65^. The transformed *N. benthamiana* tissues were imaged two days after infiltration using a laser scanning confocal microscope (Leica SP8). For each experiment, at least three independent infiltrations were performed. Images were taken in multi-track mode with line switching when multifluorescence was used. For GFP/RFP or FITC/TRITC combination, samples were excited at 488 and 552 nm and detected at 510–550 and 590–650 nm, respectively. For CFP/GFP/RFP or GFP/YFP/mCherry combination, CFP or GFP was excited at 458 nm and detected at 470–510 nm; GFP or YFP was excited at 514 nm and detected at 550–580 nm; RFP or mCherry was excited at 552 nm and detected at 590–650 nm. Images were taken using a 63x oil immersion objective (NA = 1.4), with scan speed of 400 Hz at a resolution of 1024 × 1024 pixels. To capture the dynamics of TraB, VAP27, mitochondria and autophagosomes in Arabidopsis, cells from the root elongation zone and cotyledon were acquired at a fixed rate of 0.876 s per time point for 1 min. Airy scanning super-resolution confocal imaging was performed using the Zeiss LSM 880 confocal laser scanning microscope in fast AiryScan mode, and using a Zeiss C PL APO x63 oil-immersion objective lens (NA = 1.4). Airy scanning confocal image raw data was processed using the Airyscan Processing tool on Zeiss Zen Black software. FRET-FLIM was performed as previously described^18^. For BiFC analysis, full-length CDS of TraB1a and VAP27-1 were sub-cloned into pCL112-nYFP and pCL113-cYFP plasmids, respectively^66^. Transiently transformed *N. benthamiana* were imaged two days after infiltration using a Leica SP8 confocal microscope. Images were excited at 514 nm and detected at 550–580 nm for YFP. For TMRM staining, seedlings were grown on MS medium for 5 days, and transferred into MS liquid medium with 500 nM TMRM for 10 min. Samples were excited at 552 nm and detected at 590–640 nm. Mitochondrial aspect ratio and fluorescence intensity was measured using Image J.

### Phylogenetic analysis

To identify TraB family homologues, the predicted proteins of each genome were searched using BLASTP with *Arabidopsis thaliana* TraBs as input sequences. Used databases were NCBI (https://blast.ncbi.nlm.nih.gov/Blast.cgi), Phytozome (https://phytozome.jgi.doe.gov/pz/portal.html), and Orange Genome Annotation Project (http://citrus.hzau.edu.cn/cgi-bin/orange/search). Multiple alignments were constructed with the TraB domain amino acid sequence through SMART database (http://smart.embl-heidelberg.de). The TraB family tree was generated from multiple alignments by applying the Maximum Likelihood method based on the JTT matrix-based model^67^ to a bootstrapped dataset with 1000 replicates. Initial tree(s) for the heuristic search were obtained automatically by applying Neighbor-Join and BioNJ algorithms to a matrix of pairwise distances estimated using a JTT model, and then selecting the topology with superior log likelihood value. The tree is drawn to scale, with branch lengths measured in the number of substitutions per site. The analysis involved 44 amino acid sequences. All positions containing gaps and missing data were eliminated. There was a total of 74 positions in the final dataset. Evolutionary analyses were conducted in MEGA7^68^ v1.0.5877. See Supplementary Sequence data for a complete list of all protein sequences.

### Arabidopsis phenotype studies

For general growth phenotype studies, seedlings were grown on half MS medium for 7 days, and transferred to soil ina growth chamber (22 °C, 70% relative humidity, 14 h light/10 h dark cycle). For DNP treatment and recovery, Arabidopsis seedlings were grown on MS medium for 5 days, and transferred onto medium supplemented with either DMSO or 50 μM DNP for 4 h, then the seedlings were transferred onto MS medium to recover for 2 days. Images were captured using a digital camera (Canon, EOS 80D), and the root length was measured using Image J.

### Yeast-two-Hybrid

Testing and control combinations of plasmids were co-transformed into yeast AH109 strain using the Frozen-EZ Yeast Transformation II Kit (ZYMO, T2001) and cultured on a SD/-Leu-Trp selective plate at 30 °C for 2–4 days. Three independent colonies of each combination of plasmids were picked and dotted on a SD/-Leu-His-Trp selective plate with 5 mM 3-AT. Combinations of pGBKT7-53 + pGADT7-T and pGBKT7-lam + pGADT7-T were used as positive and negative controls, respectively.

### Proteomics screen, co-immunoprecipitation and western blotting

The TAP-tag purification and mass spectrometry was performed by a proteomic service performed in Ghent University, as described previously^69^. The immunoprecipitation assays were performed using GFP-Trap_A (Chromotek, gtma-20) as described previously^6^. For detection, the membrane was incubated in 2xTBST buffer with 5% milk prior to primary antibody incubation (1:500 for anti-VAP27; 1:2000 for anti-GFP, Biorbyt orb323045) at room temperature for 3 h. After three washes in TBST buffer, the membrane was probed with HRP conjugated mouse secondary antibody (Yeasen, 33201ES60) at 1:5000 and developed using a Super ECL reagent (Yeasen, 36208ES60).

### Electron Microscopy

All samples were prefixed in 2.5 % glutaraldehyde (v/v in 0.1 M phosphate buffer, pH 7.2) for 2 h, and then rinsed 3 times with 0.1 M phosphate buffer (pH 7.2). They were post-fixed in 1 % OsO_4_ for 2 h, followed by three 15 min rinses with phosphate buffer. Afterwards, the samples were dehydrated through an acetone series (30 %, 50 %, 70 %, 90 %, 100 %, 100 %, 100 %) (v/v in dd H_2_O) at room temperature, samples were incubated for 20 min at each concentration. Then the samples were infiltrated in a graded scale of 3:1, 1:1, 1:3 (v/v) acetone/SPI-PON 812 resin and, as the last step, in 100 % (v/v) SPI-PON 812 resin (SPI Supplies, West Chester), for 12 h per step. Samples were embedded in SPI-PON 812 resin and polymerized at 60 °C for 48 h. Ultrathin sections (80 nm) were prepared using an EM UC7 Ultracut ultramicrotome (Leica, UC7). Sections were observed and photographed using a transmission electron microscope (Hitachi H-7650) at an accelerating voltage of 80.0 kV. The samples for immunogold labelling were fixed and probed with a VAP27 antibody as describe previously^19^.

### Drug treatment and autophagy assays

For Conc A treatment, Arabidopsis seedlings were treated with either DMSO or 1 μM Conc A for 8–12 h prior to imaging. For the Conc A & DNP treatment, Arabidopsis seedlings were first grown on MS medium for 5 days, and transferred into liquid MS medium supplemented with Conc A for 12 h, then DNP was added into the same medium to a concentration of 50 μM, the seedlings were incubated for an additional 2 h prior to microscopy.

For autophagy-flux tests, seedlings were treated with 1 μM Conc A before western blotting, the dilution ratio of primary antibody was 1:2000 for anti-GFP (Biorbyt orb323045) and HRP conjugated mouse secondary antibody (Yeasen, 33201ES60) was 1:5000.

### Real-time qPCR

RNA from various samples (e.g. roots, stems, leaves, flowers) were extracted according to the instructions of the Universal Plant RNA Isolation Kit (NOBELAB, RNE35). The concentration of RNA was measured by NANODROP 2000 ultraviolet. 1 μg RNA was used to synthesize the first-strand cDNA by HiScript II Q RT SuperMix (Vazyme, R223). Finally, the gene expression of *AtTraB1a* and *AtTraB1b* were detected in a qPCR system (Roche, Lightcycler 480). ChamQ Universal SYBR qPCR Master Mix (Vazyme, Q711) was used to perform the reaction. The relative expression level of each gene was quantified with the comparative threshold cycle method, using actin as the internal reference. Reactions for each of the three biological replicates were performed in duplicate. The primer sequences used in RT-qPCR are shown in Supplementary Table S1.

### Starch Assay

Rosettes of four-five weeks old plants were collected and immediately transferred to 95% ethanol, and boiled until completely decoloured (about 10–15 mins). Samples were stained in 5% Lugol’s solution (5% [w/v] I_2_ and 10% [w/v] KI) for 10 min, and then destained in water until a clear background was obtained before imaging^70^. For starch content measurement, lugol-stained samples were washed using deionised water until decoloured, air dried at room temperature and ground into powder. The powdered tissue was washed three times with 2 ml of diethyl ether and subsequently washed twice with 80% ethanol, and then twice with ddH_2_O. The washed powder was boiled in 5 mL ddH_2_O for 15 min. 980 μL of the supernatant was mixed with 20 μL of staining solution [5% (w/v) I_2_ and 2% (w/v) KI]. This reaction was subjected to colorimetric determination at 660 nm. The standard curve was performed with standard starch (Sigma, 33615).

### ATP measurement

ATP, ADP, and AMP content was measured using HPLC. Sample preparations and adenylate standards for the measurement were performed as previously reported^71^. The analysis of adenosines was performed by HPLC on a C_18_ column (Acclaim^®^ PolarAdvantage II, 4.6 × 150 mm, 3 μm, 120 Å) connected to an HPLC system (1525 binary HPLC pump coupled with 2998 photodiode array detector and 2707 autosampler, Waters). The HPLC analysis was carried out as described previously^72^. The gradient for separation of adenosine derivatives was optimized as follows: 0 min, 100% A; 2 min, 95% A and 5% B; 6 min, 80% A and 20% B; 7 min, 75% A and 25% B; 8–10 min, 100% A (running buffer A: 0.06 M K_2_HPO_4_, 0.04 M KH_2_PO_4_, pH 7.0; running buffer B: 100% Acetonitrile). The flowrate is set to 1.2 ml/min.

### Statistical analysis

All statistical graphs were performed using the GraphPad Prism software (ver. 7.00). The results were compared using the Student’s t tests and data were expressed as the mean ± SEM (Std. Error of Mean) with each symbol. P value below 0.05 was considered significant.

### Accession numbers

The Arabidopsis Genome Initiative locus identifiers for the genes mentioned in this article are TraB1a (AT1G05270), TraB1b (AT2G32340), TraB2 (AT5G52030), VAP27-1 (AT3G60600), VAP27-3 (AT2G45140), VAP27-4 (AT5G47180), VAP27-6 (AT4G00170), ATG8a (AT4G21980), ATG8e (AT2G45170), ATG5 (AT5G17290), and ATG7 (AT5G45900).

## Acknowledgements

We thank Xiaolu Qu and the microscopy core facilities of the College of Horticulture & Forestry sciences; Jianbo Cao and the Public Laboratory of Electron Microscopy, Huazhong Agricultural University. We thank Ruixi Li (South China Technology University) for providing the transgenic Arabidopsis line expressing Mito-YFP. We thank Jinli Gong, Meiyan Shi and Jingze Zang (Huazhong Agricultural University) and Christine Richardson (Durham University) for their help in this work. The project was supported by NSFC grants (no. 91854102; 31772281), HZAU Scientific & Technological Self-innovation Foundation (2017RC004) to P.W; and a BBSRC grant (BB/G006334/1) to P.J.H.

## Author Contribution

P.W. and P.J.H. conceived and supervised the project, C.L. and P.D. performed most of the experiments, C.L., P.D., P.J.H. and P.W. wrote the manuscript. T.Z. and J.K. generated the CRISPR mutant. Y.F. performed the phylogenetic studies. X.L, and Y.C helped on ATP analysis and HPLC studies. G.D.J performed the TAP-tag MS screen. T.Z., and Y.F. helped with confocal imaging, material preparation and data interpretation.

## Data availability

The authors declare that all data supporting the findings of this study are available within the article and its Supplementary Information files, or from the corresponding author upon reasonable request.

## Competing Interests

The authors declare no competing interests.

## Notes

### Competing Interest Statement

The authors have declared no competing interest.

